# Leveraging Neural Network Models for Drug Repurposing: A case study on Cardiac Hypertrophy

**DOI:** 10.1101/2025.01.28.635194

**Authors:** Rasmus Magnusson, Markus Johansson, Sepideh Hagvall, Jane Synnergren

**Affiliations:** Systems Biology Research Center, School of Bioscience, University of Skövde, Skövde, Sweden; AstraZeneca R&D, Chief Medical Office, Global Patient Safety, Pepparedsleden 1, Mölndal, Sweden; Gothenburg University, Department of Molecular and Clinical Medicine, Institute of Medicine, Sahlgrenska Academy, Gothenburg, Sweden

## Abstract

**Background:** Drug repurposing has emerged as an attractive strategy in contemporary pharmaceutical research, presenting an opportunity to expedite drug discovery, minimize developmental costs, and mitigate risks associated with developing new pharmaceuticals. In this study, we investigated a novel approach based on deep learning of human transcriptomic mechanisms for systematic identification of additional therapeutic potential in preexisting drugs.

**Method:** We trained a composite feedforward neural network model using gene expression data sourced from the ARCHS4 compilation of the GEO, encompassing extensive human datasets. Subsequently, disease-associated gene expression data were generated from our stem cell-derived *in vitro* model of cardiac hypertrophy induced by Endothelin-1 stimulation. These data were employed to identify latent variables associated with genes showing differential expression due to Endothelin-1 stimulation. By examining the differential expression profiles within the model’s latent space, we successfully correlated the disease signal with known drug targets found in pharmaceutical compounds cataloged in DrugBank.

**Results:** The model accurately encoded additional disease-related genes beyond the curated gene set, demonstrating its ability to generalize disease associations. Leveraging the model, we identified potential drug candidates, such as lapatinib and amiodarone showing promise in mitigating proBNP concentration associated with cardiac hypertrophy.

**Conclusion:** This study demonstrates the power of deep learning of human transcriptomic mechanisms in swiftly identifying new therapeutic potentials for existing drugs, highlighting the pivotal role of artificial intelligence technologies in accelerating drug development for other complex medical conditions.

**Highlights:**

- An interpretable neural network model for identification of candidate drugs for drug repurposing
- Encoding expression data from our cardiac hypertrophy model highlights important disease mechanisms
- Lapatinib and amiodarone experimentally validated as candidate drugs for cardiac hypertrophy therapies

## 1. Introduction

Drug repurposing involves utilizing approved drugs and failed candidates to target new indications, thereby identifying alternative therapeutic uses for existing medications initially developed for different medical conditions. Exploring existing drugs for their potential therapeutic effects on various diseases or conditions proves economically beneficial [1], rather than developing entirely new drugs. This process typically integrates advanced computational analysis, high-throughput analyses, and experimental testing to identify potential candidates, particularly those demonstrating promise in targeting molecular pathways associated with new diseases or conditions.

While repurposing drugs for new medical conditions necessitates extensive preclinical and clinical testing to confirm efficacy and safety, it offers substantial advantages compared with *de novo* drug development [2]. This approach facilitates significantly shorter development time as time consuming early steps in the standard drug development process can be omitted or shortened (Fig. 1). For example, since these drugs have already undergone prior safety and efficacy testing, they bypass several regulatory and safety requirements and prove to be more cost-effective [2]. Repurposed drugs also carry lower risks due to established safety profiles, minimizing unforeseen adverse effects compared to entirely new compounds. These benefits lead to significantly reduced costs and provide possibilities for novel therapies for medical conditions that currently lack efficient treatments. Furthermore, drug repurposing presents opportunities for combined therapies, fostering synergistic treatments for complex diseases [3–6].

**Figure 1:**
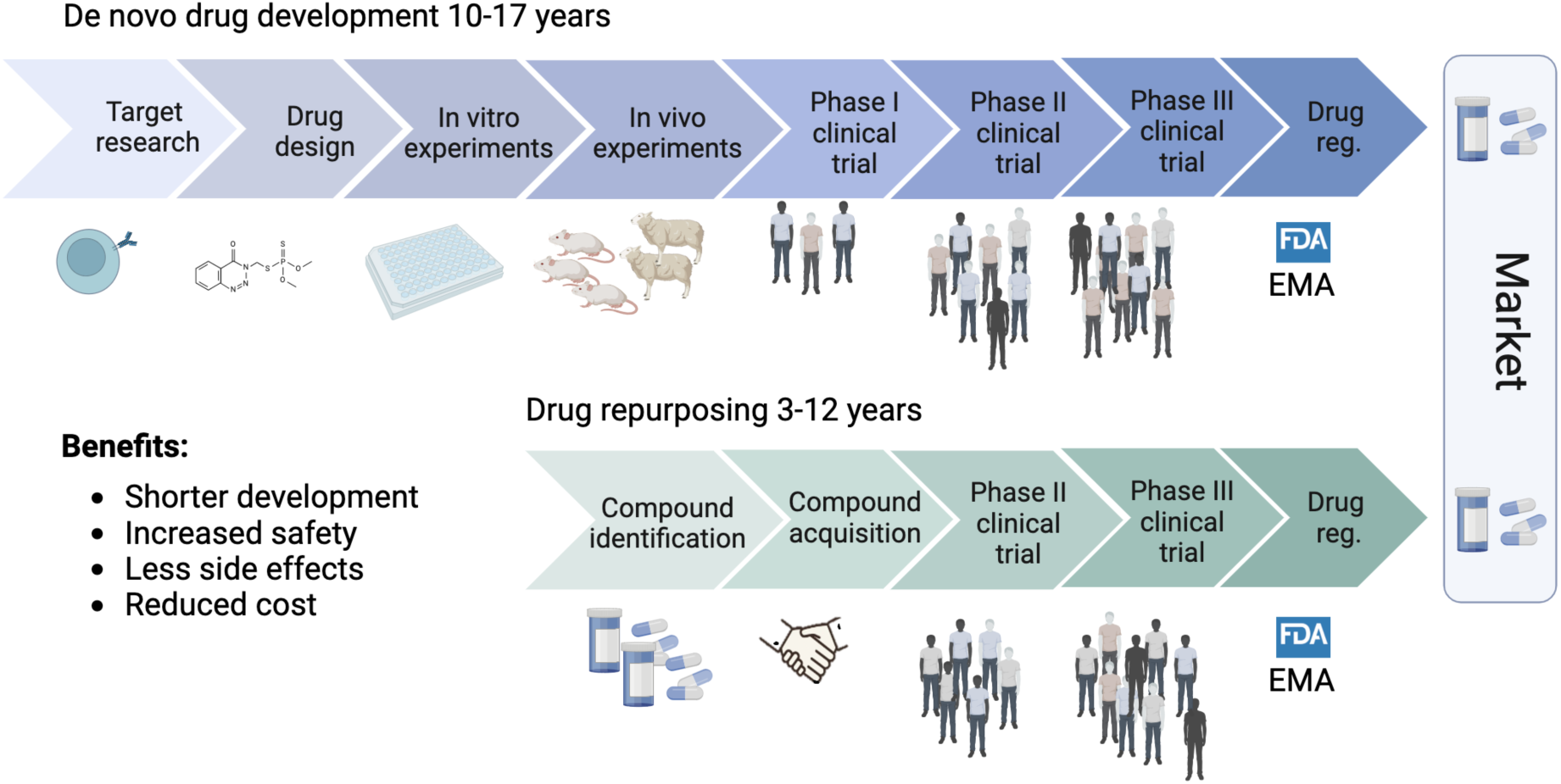
Overview of de novo drug development and the main steps performed during the traditional drug development process that typically takes between 10-17 years. In green are the main steps of drug development based on drug repurposing outlined, highlighting shorter development, increased safety, less side effects, and reduced cost.

Another important aspect of drug repurposing is the potential for accelerated development of therapies for novel diseases, which is a huge advantage when specific drugs are not yet available or developed. The unpredictable emergence of infectious diseases and the lack of available treatments for rare diseases pose significant global health challenges [7]. The most recent example is the COVID-19 pandemic, which affected many millions of individuals worldwide. During this time, medical practitioners were struggling to find a rapid cure for hospitalized patients to improve the conditions of the patients and minimize outbreak severity. As drug repurposing matches the urgency of finding effective treatments it was applied as an effective strategy in the search for more efficient treatments for COVID-19 [8]. One such example is baricitinib, an anti-inflammatory drug approved for rheumatoid arthritis, which was repurposed and authorized for emergency COVID-19 treatment based on encouraging trial results [9].

Different strategies are described in the literature to perform drug repurposing [10–14], and these can be classified into *experimental*- or *computational* approaches. In the experimental approaches binding assays and phenotypic screenings are applied to identify binding interactions of ligands to assay components and identify lead compounds from large compound libraries. The computational approaches include drug centric strategies, target-, network/pathway-, AI-, knowledge-, and signature-based approaches. Importantly, these computational methods accelerate the drug discovery process by effectively utilizing information such as cheminformatics, bioinformatics, network biology and systems biology [11]. In the study presented here we applied a combination of target-, pathway-, and AI-based strategies for drug repurposing (Fig. 2), which demonstrated to be successful in terms of identification of existing drugs for novel indications of cardiac hypertrophy. Based on our result we propose an extension of the framework described by Parvathaneni [10], to also include AI based approaches, as highlighted in Fig. 2.

**Figure 2:**
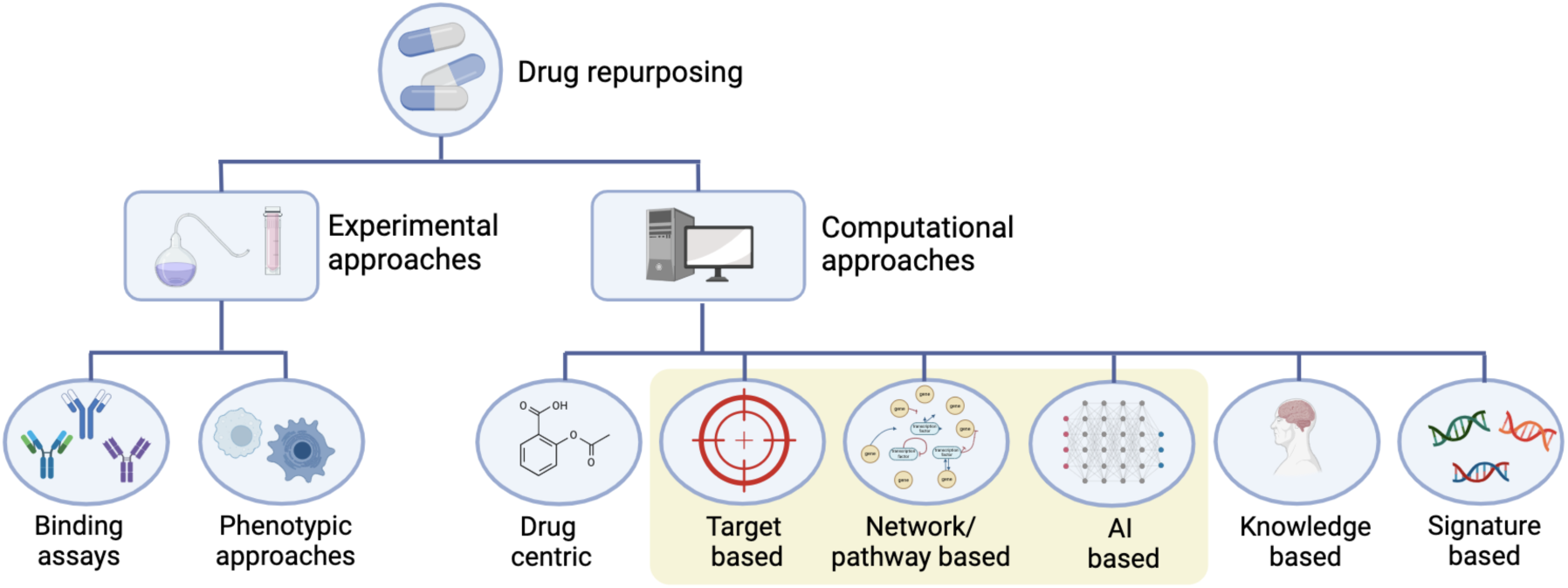
A framework of strategies to perform drug repurposing inspired by Parvathaneni et al. [10]. Both binding assays and phenotypic screening approaches are applied to identify binding interactions of ligands, and identify lead compounds from large compound libraries, and these are classified as *<u>experimental approaches</u>* [6]. The *<u>computational approaches</u>* are generally high-throughput, and thus efficient tools in discovering therapeutic agents for novel indications.

The process of matching known drugs to new diseases is however not straight-forward. There are several reasons for this, including that diseases often involve one or several mechanisms that are seldom fully understood, and that these mechanisms induce changes across the interactome. Nevertheless, the usage of computational biology methods has proven to be an important tool to predict drug repurposing candidates, with numerous success cases [12–14]. This development has recently become of interest from an artificial intelligence and artificial neural network (ANN) modeling perspective [15]. An interesting work by Kuenzi et al. (2020) [16] presents an ANN framework named DrugCell for repurposing drugs. Notable, this framework was constructed by incorporating 2,086 known biological processes from the Gene Ontology database into the structure of the latent space, with the rationale to build an interpretable model structure.

In this study we redesigned the approach used by Kuenzi et al., by building an ANN model with known disease gene sets coded into the latent space. In detail, we used groups of curated gene associations of 1,336 human diseases in the DisGeNET database [17] to build a composite neural network with a latent space of 1,336 variables, each corresponding to a unique disease set.

This novel drug repurposing approach was then applied to a case study of cardiac hypertrophy to evaluate the performance. Cardiac hypertrophy is a physiological or pathological condition characterized by an increase in the size of the heart muscle, specifically the myocardium[18]. The physiological hypertrophy occurs as a normal and adaptive response to increased workload on the heart, such as during regular exercise [19] or pregnancy [20]. It leads to an increase in the size of individual cardiomyocytes without significant adverse effects on the heart function. On the other hand, pathological hypertrophy is often associated with chronic stress on the heart and is generally detrimental to the heart function [21]. The heart muscle becomes thicker and stiffer, which impair its ability to contract effectively and pump the blood in the circulatory system [22]. This condition is often associated with detrimental changes in gene expression, signaling pathways, and structural remodeling of the heart. It can develop to a serious maladaptive response that can lead to heart failure, arrhythmias, and other cardiovascular complications. Common causes to pathological hypertrophy include hypertension, aortic stenosis, and cardiomyopathies [23–26]. Notably, even if the cause of the pathological hypertrophy is removed, by the use of existing heart failure therapies, the condition is often retained and most patients do not recover from its diseased condition [27]. Novel therapeutic approaches are needed to help patients to recover and protect them from future cardiac failure.

This study presents a novel approach to drug repurposing through the integration of artificial intelligence methodologies and transcriptomic high-throughput data. By specifically embedding disease-associated gene sets into a composite neural network structure, we demonstrate a capacity to decipher intricate transcriptomic signatures underlying various diseases, with cardiac hypertrophy as a focal case study. This work illuminates the power of neural network learning in comprehending disease mechanisms, offering a transformative approach to swiftly identify therapeutic avenues for complex medical conditions.

## 2. Material and methods

### 2.1. Transcriptomics Data Curation

Utilizing gene expression data sourced from the extensive compilation of human RNA-seq experiments in the ARCHS4 database [28], we initially filtered out pseudo-genes and performed a natural log transformation on the counts of the remaining 27,486 unique genes. To eliminate low-quality entries, such as when very few counts were present in a sample, we excluded samples where over half of the genes showed no detectable expressions. Consequently, we obtained 27,627 curated gene expression profiles, and reserved 8,687 of these for model testing.

### 2.2. Data analysis and model development

Herein, we used a feed-forward neural network (FFNN) implemented in Keras. The model’s architecture aimed to compress gene expression data into a specifically designed latent space embedding disease-related information. Leveraging the DisGeNET database [17], we constructed 1,336 individual fully connected feed-forward neural networks, each uniquely representing a disease from DisGeNET. As the DisGeNET includes numerous diseases with a limited set of associated genes, we opted to only include the 1,336 cases with 10 or more associated genes to increase the validity of our results. These disease-specific models were constructed to take the associated gene expressions as input. Inspired by the model design presented in our recently published paper de Weerd et al. 2024 [29], the model featured two hidden layers with 20 nodes each, and culminated in a single node as output. Subsequently, these 1,336 outputs were passed to a fully connected hidden layer comprising 250 nodes that were connected to an output layer representing the gene expression levels of the entire human transcriptome (Fig. 3A).

**Figure 3.**
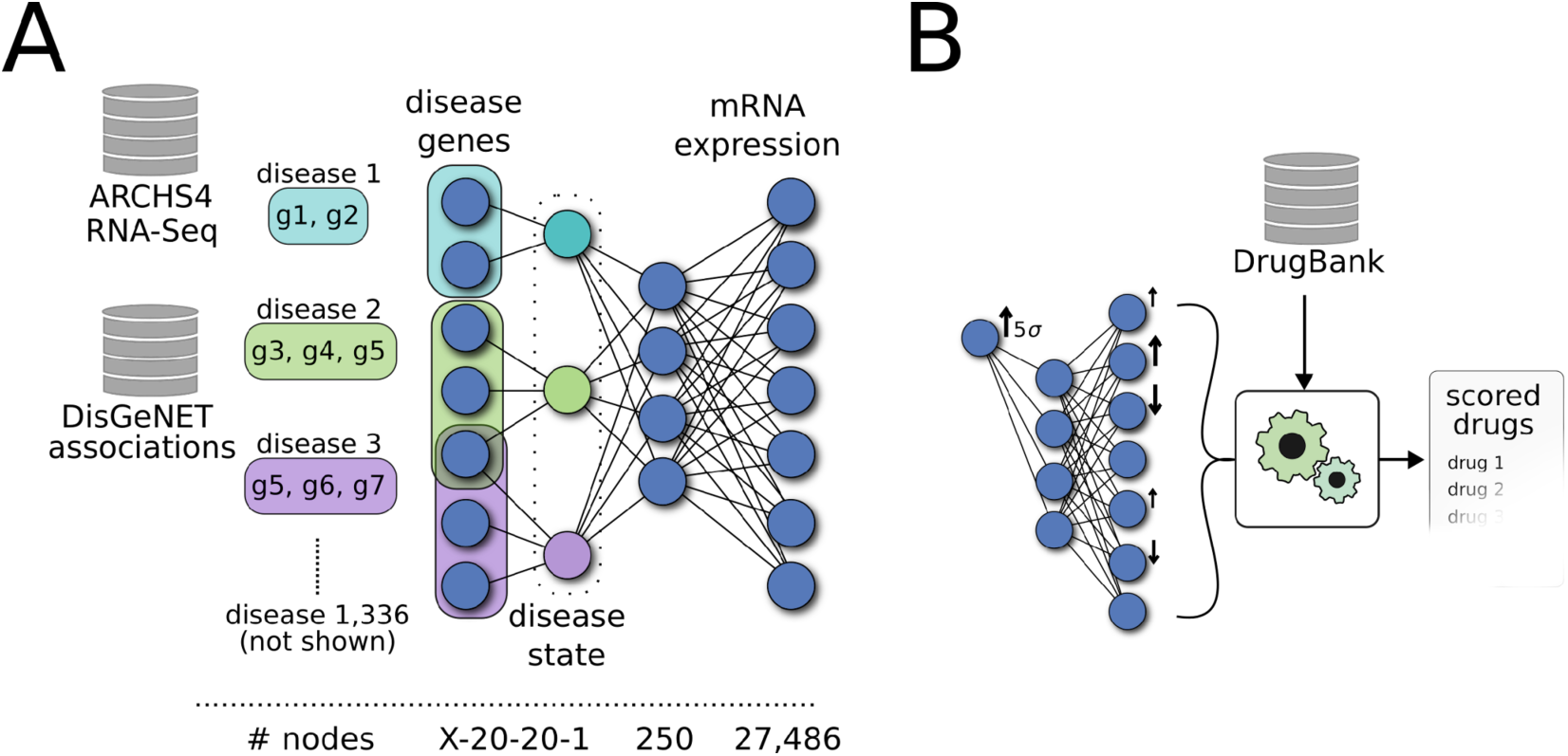
Model design and usage. A) We built a feed-forward neural network regression model, using gene expression profiles from the ARCHS4 dataset as input and output. We aimed to embed an explainable latent space, with a layer representing known diseases. To this end, we used gene-disease associations from the DisGeNET, and built 1,336 independent sub-models, i.e. one per disease, all acting as input to the latent space. For a disease with X associated genes, the sub-model would have X inputs, fed to two subsequent layers with 20 nodes each, and finally one single output node. The full model was trained to predict the gene expression of all 27,486 genes. B) We perturbed each node in the most compressing hidden layer to associate disease with decoder genes. The decoder output was observed and matched to the target genes of known drugs.

We used a principal component analysis (PCA) to extract features from the latent space encodings of the model. As such, we used the 8,687 test data profiles and calculated the components in the resulting 8,687×1,336 matrix. In order to extract a stronger signal, we chose to independently amplify the data ten-fold along each of the 1,336 components. Next, we aimed to study which genes were encoded on each component. Since each component amplification resulted in a continuum of gene associations, we opted to study the respective top-changing 100 genes. In our analysis of the biological processes encoded on each principal component, we individually mapped the 100 genes of each component to a protein-protein interaction (PPI) network. Specifically, we used the STRING and the FunCoup PPI databases, respectively. In gene-gene distance calculations, we used the harmonic mean distance, as this metric is applicable even in the case where two genes are unconnected on a PPI, and the distance is annotated as infinite.

Inspired by the DrugCell method developed by Kuenzi et al. in 2020 [16], this model design facilitated an interpretable latent layer, where each of the 1,336 hidden nodes encoded the gene expression associated with a particular disease. We used the same activation function across all nodes, employing the exponential linear unit (elu), and optimized the models to minimize mean squared error using the Adam algorithm [30]. We trained the model to convergence using batches from the 27,627 gene expression profiles in the training dataset. The model was trained using CPU resources supplied by National Academic Infrastructure for Supercomputing in Sweden (NAISS), consuming <1000 core hours. The model and accompanying code can be downloaded at https://github.com/rasma774/composite_disease_model.

### 2.3. Drug repurposing candidate inference from case-control data

We conducted an analysis using RNA-seq data obtained from our previous study involving induced pluripotent stem cell (iPSC)-derived cardiomyocytes, stimulated with ET-1 to induce cardiac hypertrophy, for which the data is publicly available [31]. This dataset served as the basis for detecting significant differences in activations within the hidden layer of our model. We Z-score transformed the activation of each hidden node and selected the node with the highest differential activation as an effect of the ET-1 treatment for further study. In other words, we chose to study the embeddings of the node with the highest relation to ET-1 stimulation. To analyze what genes in the output layer were associated with this hidden node, we employed a 5-standard-deviation activation increase, where the standard deviation was calculated from the validation data in ARCHS4 (Fig. 3B).

These disease-associated genes were then cross-referenced with pharmaceutical compound targets, as referenced in Guney et. al [32], to rank the drugs. We utilized the gene responses from the 5-standard deviation activation such that we, for each drug, compared the average response in the known drug targets to that of all genes in the decoder. The rationale is that if the drug is associated to the most differentially activated node in the ET-1 dataset, the drug’s target genes should respond to such a perturbation.

### 2.4. Analysis of drug properties and characteristics

The list of ranked candidate substances was further manually filtered and substances incompatible with in vitro cell culturing due to limited solubility, high toxicity etc. were excluded. Then, the substances’ predicted effects on the cardiac hypertrophy signaling pathway were investigated. Only the substances that showed inhibition of this signaling pathway and a reduction in a cardiac hypertrophy outcome were considered. The highest-ranked drugs based on their calculated score were selected, one per drug target class, for experimental validation.

The selected drugs include Amiodarone, Binimetinib, Bosentan, Disulfiram, Lapatinib, and Regorafenib. Amiodarone is an antiarrhythmic drug used to treat and prevent conditions such as ventricular tachycardia and atrial fibrillation. Binimetinib is a selective mitogen-activated protein kinase 1/2 inhibitor used to treat a variant of metastatic melanoma. Bosentan, an endothelin receptor antagonist, is used to treat pulmonary arterial hypertension. Disulfiram aids in the treatment of chronic alcoholism by inducing an acute sensitivity to ethanol. Lapatinib, a tyrosine kinase inhibitor, is used in the treatment of breast cancer. Regorafenib, a multi-kinase inhibitor, is utilized in various forms of cancer treatment.

### 2.5. Cell culture for experimental validation

Human cardiomyocytes (CMs) derived from hiPSC line ChiPS22 were obtained from Takara Bio (Takara Bio Europe AB). The CMs were cryopreserved at day 19 following the onset of differentiation using STEM-CELLBANKER (cat 11890, Amsbio). Cryopreserved CMs were thawed in CM medium (Advanced RPMI, B27 1x, Glutamax 1x, (Thermo Fisher Scientific) supplemented with Y27632 (10 µM) (cat. Y0503, Sigma-Aldrich) and plated at 3.75*10^5^ cells/well in a 24-well plate precoated with 5 µg/cm^2^ Fibronectin solution (5 µg/cm^2^) (cat. F0895, Sigma-Aldrich). One day after thawing, the culture medium was changed to CM medium and cells were recovered for a total of 6 days before starting the experiments with Endothelin-1 (ET-1). Medium (0.4 ml/cm^2^) was changed every second day. For hypertrophy induction, ET-1 (cat. E7764, Sigma-Aldrich) was dissolved in DMSO and then added to the culture medium. 24h after ET-1 was added, culture media was changed to CM media with ET-1 + substance. Amiodarone was dissolved in methanol while the other substances were dissolved in DMSO. The cells were treated with ET-1 + substance for 24h before samples were collected. The corresponding volume of DMSO or methanol was added to the control cells in parallel. All experiments were repeated at least three times. The study design is outlined in Fig. 4.

**Figure 4.**
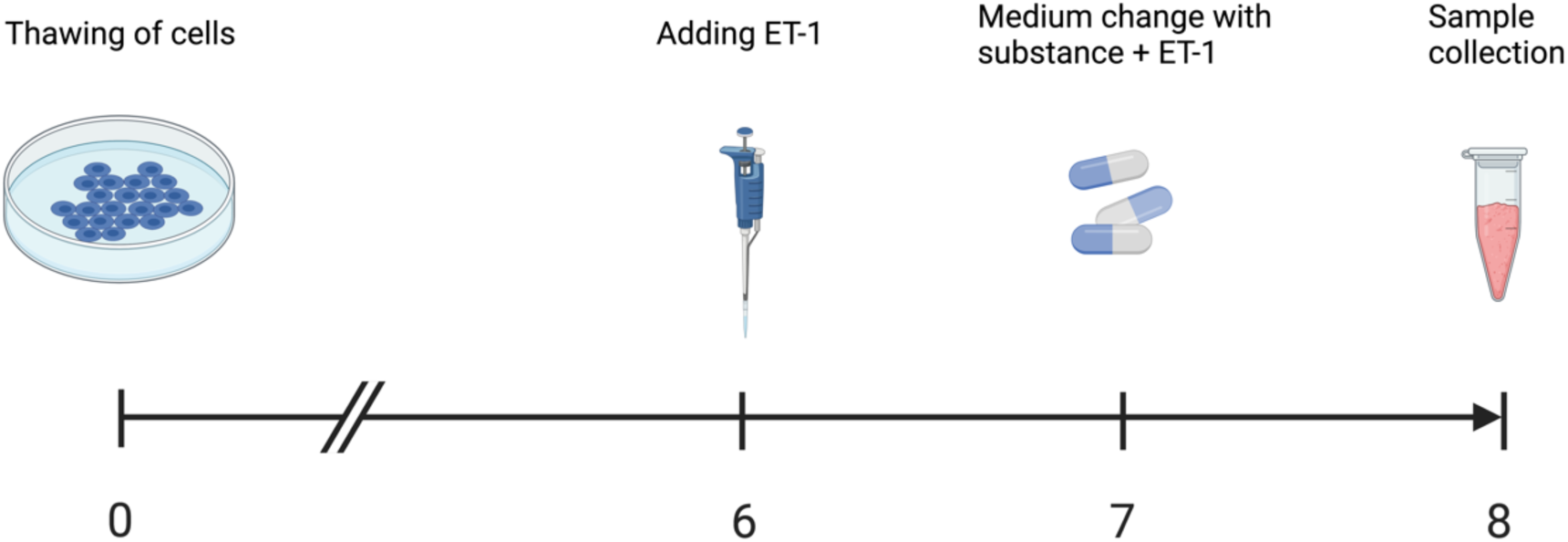
Overview of the experimental design. The hiPSC-CMs were thawed and cultured for 6 days before ET-1 was added to the cells. On day 7, the medium was changed, and a new dose of ET-1 was added. On day 8, the medium was collected and stored for later analysis.

### 2.6. Enzyme-Linked Immunosorbent Assay (ELISA)

Conditioned media was collected from all the samples 48h after ET-1 was added to the media, and subsequently 24h after the potential hypertrophy inhibitors were added. The media was centrifuged (5000x g, 5 min) and the supernatant was collected and stored at −80° C for subsequent analysis. The concentration of proBNP in the samples was analyzed using ELISA kits (EHPRONPPB, Thermo Fisher Scientific) according to the manufacturer’s instructions.

## 3. Results

Expanding upon recent advances that integrate prior biological knowledge into deep learning models, our investigation aimed to determine if incorporating disease-gene associations within a feedforward neural network (FFNN) could predict specific drug candidates. Initially, we constructed a FFNN utilizing 1,336 disease gene sets as input (Fig. 1A). Finally, leveraging the model (Fig. 5), we identified potential pharmaceutical candidate drugs, focusing on cardiac hypertrophy as a case study. Experimental validation of these results showed a significant reduction in proBNP levels in an in vitro hypertrophy disease model following treatment with amiodarone and lapatinib. The reduction was in similar levels as for the ET-1 receptor antagonist bosentan, which was included as a positive control, indicating their potential therapeutic impact.

**Figure 5:**
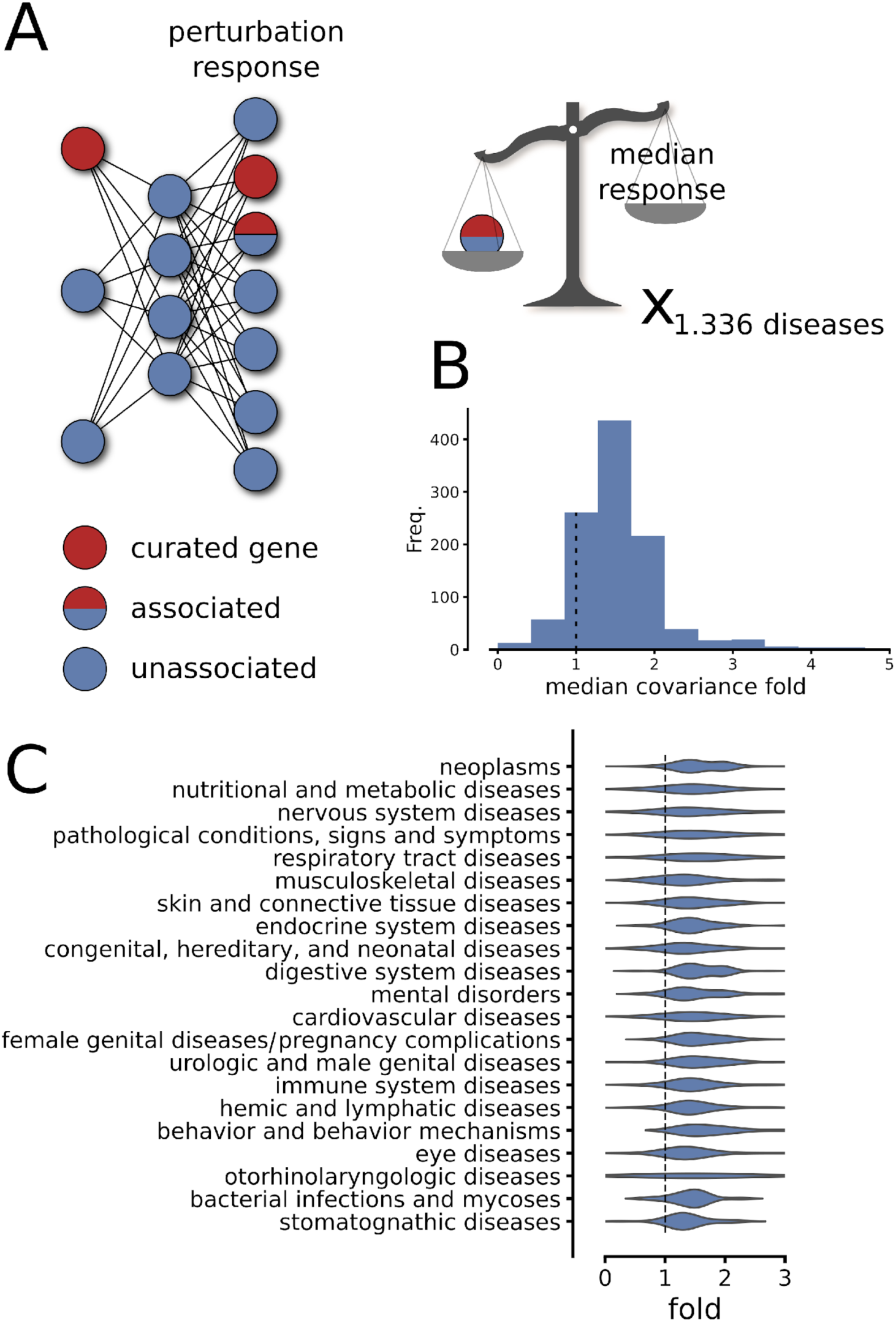
The model training embedded relevant disease structures. A) To test how the model generalized in terms of capturing disease mechanisms, a set of disease-gene associations of lesser confidence and unseen by the model was used. For each disease-embedding in the latent space, we recorded how a perturbation of 5 standard deviations changed these associated genes in relation to the median change in the predicted expression. B) The median of the response of the associated genes was divided by the overall median response, which is referred to as the covariance fold. Shown is a histogram of these 1,336 folds. Note that the distribution mean is above the expected value of 1. C) By stratifying the results shown in B) into disease classes, we found a clear signal across all disease types.

### 3.1. A set of 1,336 disease-specific variables can accurately predict the transcriptome

We aimed to integrate biological a priori knowledge into our model by incorporating sets of disease-related genes for predicting the expression values of the human transcriptome. Utilizing gene-disease associations from DisGeNET, we constructed a FFNN with an interpretable latent space. Specifically, we created individual small FFNN for each gene set, compressing the expression pattern to a single variable representing a disease state. Each FFNN consisted of two hidden layers with 20 nodes each. In total, we employed 1,336 disease states and connected their outputs to a fully connected FFNN (see Fig. 3a). The entire model was optimized to predict the counts of approximately 27,000 human genes from the ARCHS4 database [28].

We tested the model’s ability to predict gene expression data by applying it to ARCHS4 gene expression profiles that were kept apart from the training data. We found the model to predict new data well, with a median Spearman’s correlation coefficient rho = 0.95. Thus, the model could accurately predict a complete gene expression profile from 1,336 hidden variables that were specifically designed to hold information of gene expressions of corresponding diseases.

### 3.2. Additional disease-associated genes became encoded in relevant latent variables

We next sought to test to what extent the model encodings had biological relevance and opted to analyze a disease association gene set that contained more genes but with lower certainty. In detail, all disease-gene associations registered in the DisGeNET but not part of the curated set that was used to design our model were downloaded and analyzed. We performed a sensitivity analysis on the decoder output with respect to each hidden node in the disease-specific hidden layer by, to each node, adding 5 standard deviations to the mean node activation, and by subsequently observing the value changes in the output layer (Fig. 5A). We found the changes of the genes that were exclusively found in the lower confidence gene-disease annotation list to have a significantly (Mann-Whitney u p < 0.05) higher value than the rest of the genes in 53% of the diseases (expected 5% under the null hypothesis, Fig. 5B-C). These results support that the model has captured relevant disease genes. Notably, the median response of those genes was higher than the expected value of 1 in 88% of all comparisons, suggesting that the number of cases where the null hypothesis is rejected would increase considerably with statistical power.

### 3.3. Extracting learned biologically relevant patterns from the model latent space

Several researchers, including our group, have previously demonstrated how FFNNs applied to transcriptomic data can learn meaningful representations of cellular biology [29, 33, 34]. To investigate the biological relevance of this model’s embedding and extract novel information, we continued our analysis by searching for biologically relevant patterns. In detail, we analyzed the principal components (PCs) of the validation data in the compressed space, such that each PC was individually amplified with a factor 10. These perturbed data were subsequently passed through the decoder, and the top 100 changing genes per PC were extracted. We hypothesized that genes that are associated to the same disease have a higher level of interaction than randomly selected genes. Thus, the internal relation between each gene set of 100 genes was then analyzed with respect to the protein-protein interaction networks in STRINGdb and in FunCoup. Results show that the top PCs correspond to gene sets that were significantly closer on the protein-protein interaction maps than what would be expected under the null hypothesis, i.e. that the genes were unrelated. In fact, the first 10 PC gene sets had a harmonic mean distance that on average was 27% (STRINGdb) and 36% (FunCoup) of the monte-carlo estimated distance of randomly sampled genes. In other words, the latent space of the FFNN appeared to have learnt known biological patterns.

### 3.4. The differential activation of hidden nodes in case-control data can be interpreted as a novel approach to disease annotation analysis

In this study, the FFNN model was specifically designed to have an interpretable latent space, with each hidden node corresponding to one disease in DisGeNET. Notably, this presents the opportunity to use the model as a gene set annotation tool, where the profiles of case-control gene expression data can be compressed and the nodes in the latent space tested for differential activation. Since each node was designed to represent a specific disease in the DisGeNET, the differential activation is interpretable as an annotation of interest.

To evaluate the model in an applied disease scenario, we utilized RNA-Seq data of endothelin-1 (ET-1) stimulated hiPSC-dervied cardiomyocytes (CMs), which we have previously demonstrated to be a model of cardiac hypertrophy [31, 35]. ET-1 stimulated samples were compared to untreated control samples, a Z-score transformation of the differentially activated nodes of the latent space was performed. When analyzing the highest ranked nodes, several relevant diseases e.g. diastolic heart failure and left ventricular hypertrophy were identified. We noted that there was a substantial degree of genes overlapping between the highest ranked terms (mean pairwise odds ratio=14.1, geometric mean of p= 0.0008), with at least one gene overlapping in all sets except the fourth top term (Parkinson disease), suggesting a homogenous disease signal.

### 3.5. Matching hypertrophy-induced differential activation in the latent space to known drug targets reveals new avenues for existing drugs

To investigate if the cardiac hypertrophy dysregulation patterns learned in the latent space could be used to predict candidates for drug repurposing, the top differentially activated node activation was increased with a factor of 5 standard deviations, as calculated in the validation data, and the changes in the output layer of the decoder were recorded. These gene changes were matched against known drug targets, as presented in [32] and a ranked list of drug repurposing candidates was extracted (Suppl table 1).

The ranked list of drugs does not indicate whether they are predicted to alleviate or exacerbate cardiac hypertrophy. It also does not indicate whether the drugs are suitable for treating cardiac hypertrophy. For example, nitrate compounds, such as nitrous acid and pentaerythritol tetranitrate, received the highest score. However, this type of substance may be unsuitable for treatment of cardiac hypertrophy as it can increase outflow tract obstruction by decreasing preload and ventricular volumes [36]. Histamine receptor agonists, such as betahistidine and histamine also showed high scoring. Analysis of these substances in Ingenuity Pathway Analysis (IPA) showed that they had a reverse predicted effect on hypertrophy, rendering them inadequate as a treatment alternative. Beta-blockers were also found to have relatively high scores with Sotalol showing the top score from this drug group. However, Sotalol has been shown to worsen symptoms of decompensated heart failure [37].

To exclude inadequate drug candidates from the list of results, a manual selection was performed. Only substances with high potential to reduce the concentration of the cardiac hypertrophy biomarker, proBNP, in our developed in vitro disease model [31, 35, 38] were selected for experimental validation. The final list of six candidates that was chosen for in vitro validation is shown in Table 1.

| Drug | Score | Target | Notes |
| --- | --- | --- | --- |
| *Bosentan* | 3,0 | ET receptor antagonist |  |
| *Lapatinib* | 2,9 | Tyrosine kinase inhibitor | Cancer drug |
| *Disulfiram* | 2,5 | Alcohol dehydrogenase inhibitor, Dopamine beta-gyrdoxylase inhibitor | Treatment for chronic alcohol abuse |
| *Amiodarone* | 2,3 | Potassium and sodium channel inhibitor | Anti-arrhythmic |
| *Regorafenib* | 1,5 | Kinase inhibitor | Cancer drug |
| *Binimetinib* | 0,6 | MEK1/2 inhibitor | Cancer drug |

Note:

Figure 6 presents an overview of the cardiac hypertrophy signaling pathway and how it is influenced by ET-1, amiodarone, lapatinib, and bosentan. While ET-1 was predicted to increase the hypertrophy response, as expected, both lapatinib and amiodarone were predicted to reduce it.

**Figure 6.**
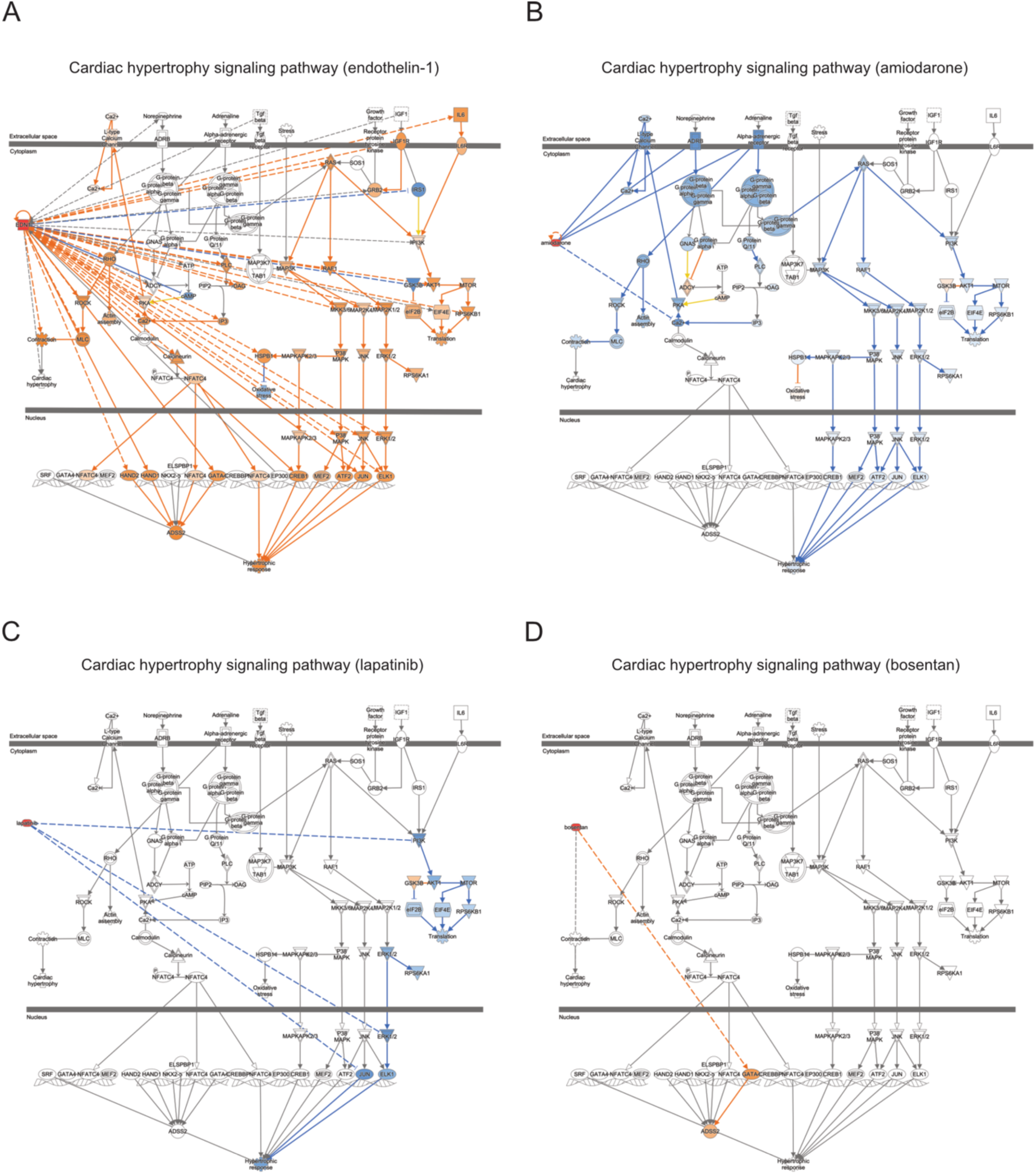
Cardiac hypertrophy signaling pathway. An overview of the signaling pathways that are involved in cardiac hypertrophy and how A) endothelin-1, B) amiodarone, C) lapatinib, and D) bosentan affect the signaling. Orange color indicates an increase in expression, and blue color indicates a decrease in expression.

### 3.6. ProBNP protein concentration after treatment with candidate drugs

The concentration of proBNP, a well-known biomarker for hypertrophy, was analyzed in our established in vitro model of hypertrophy to investigate the potential of the selected candidate drugs to reduce its concentration. Fig 4 shows an overview of the experimental validation setup. ET-1 stimulation of the CMs for 48h significantly increased the concentration of proBNP in the media for both DMSO (FC = 10.2, p = 0.0002) and methanol controls (FC = 7.8, p = 0.007) (Fig 7 a-b). ET-1 stimulation for 24h followed by ET-1 stimulation and Lapatinib treatment (1.0 and 10.0nM) for 24h significantly reduced proBNP concentration compared to only ET-1 treatment. At 5nM there was a decrease but did not reach statistical significance due to three instead of four replicates for 5.0nM (p = 0.06) (Fig.7c). Treatment with Amiodarone decreased the proBNP concentration in a dose-dependent manner. At 5.0 and 10.0nM the reduction in proBNP concentration was over 100ng/ml (Fig. 7d). The ET-1 receptor antagonist, Bosentan, significantly reduced the proBNP concentration at 10.0 and 100nM (62ng/ml and 85ng/ml respectively) but did not reach the level of reduction as Amiodarone did (Fig.7e).

**Figure 7.**
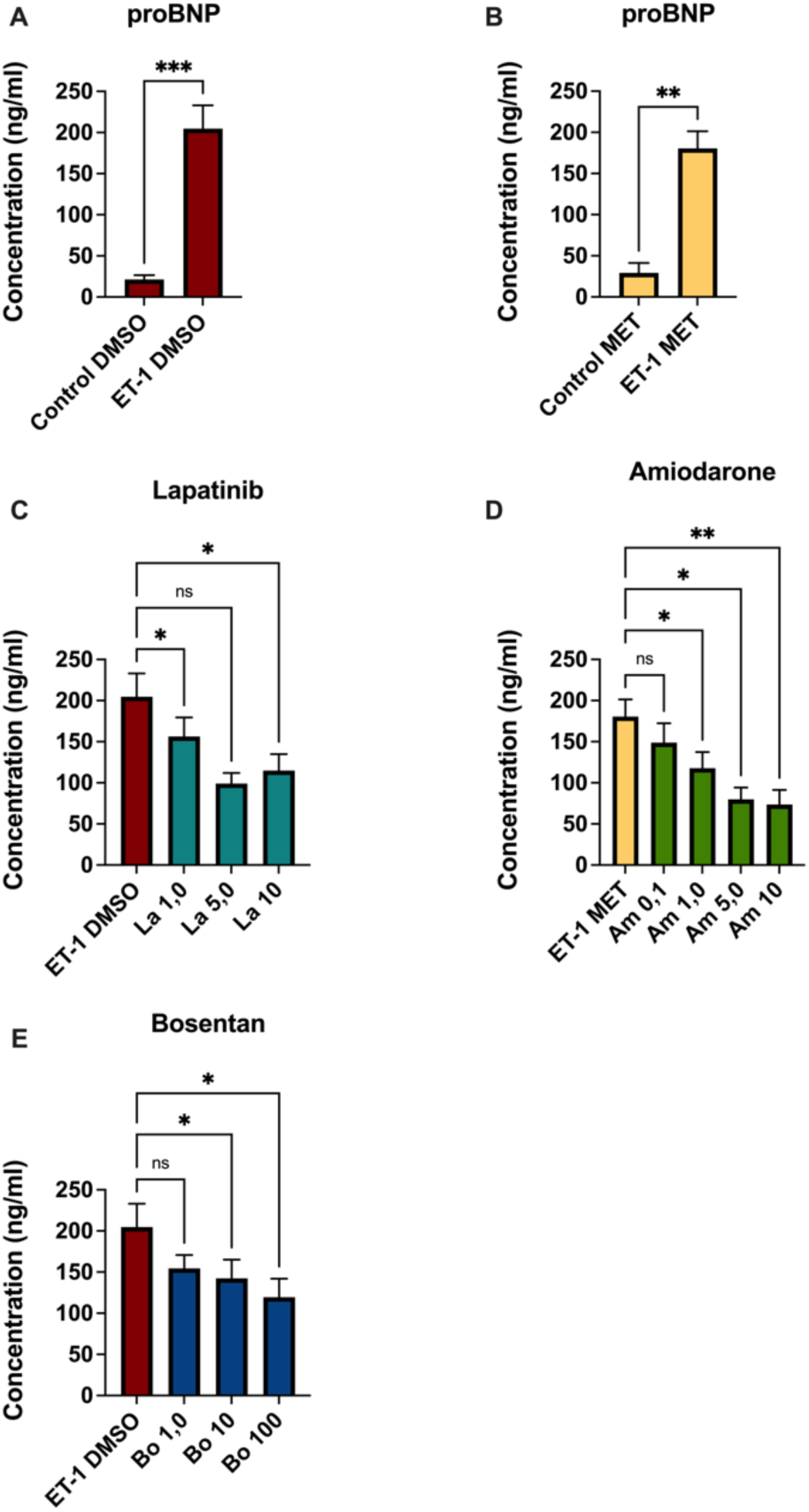
proBNP concentration analysis. Graphs showing the concentration of proBNP in the culture media, concentration in ng/ml, A) Control vs ET-1 (DMSO), B) Control vs ET-1 (Methanol), C) ET-1 (DMSO) vs Lapatinib (1,5,10uM), D) ET-1 (Methanol) vs Amiodarone (0.1, 1, 5 and 10uM) E), ET-1 (DMSO) vs Bosentan (1, 10 and 100uM). SEM is given as error bars n=4, except lapatinib 5uM (n=3) *P<0.05, **P<0.01, ***P<0.001.

Treatment with Dislufiram, Regorafinib and Binimetinib showed no significant reduction of proBNP concentration (Fig. SF1). No cytotoxic effects were observed upon treatment with ET-1 or the candidate drugs within the applied concentration range (Fig.SF2-3).

## 4. Discussion

Over the past decade the cost of drug development has surged dramatically, yet the number of new drugs approved annually has declined [39–41]. This paradox is driven by several factors, including the rising complexity of scientific research, more stringent regulatory requirements, and increased investment in advanced technologies and personalized medicine [42].

Many drug candidates fail to progress to market due to insufficient efficacy, unexpected toxicity, or adverse safety profiles observed during human trials. Despite substantial financial outlays, the approval rate for new drug candidates remains low, leading to higher costs being spread over fewer successful products. [43, 44]. Additionally, the clinical trial phase has become lengthier and more expensive due to the need for larger and more diverse patient populations to ensure safety and efficacy. These challenges have made the relevance of drug repurposing in the pharmaceutical sector progressively growing in recent years, with about one-third of approvals corresponding to repurposed drugs, some of which have even achieved blockbuster status [2, 6, 10, 43]. In other words, drug repurposing provides a promising strategy to address many challenges in traditional drug development [6, 45]. By bypassing the extensive and expensive safety testing phases of traditional drug development, there is a potential to enhance the sustainability of pharmaceutical innovation and diminish the need for extensive animal testing [42, 45, 46].

Recently, many interesting deep learning methods have been adapted for identification of drug target interactions. For example, Zhau et. al. demonstrated a convolutional network model that uses a graph convolutional network to learn the features for pairs of drug-target interactions, and with these feature representations as inputs, utilizes a deep learning neural network to classify between positive and negative drug-target interaction classes [47]. Another example is the iDrug method that unifies drug repurposing and drug-target interaction predictions into a single cohesive model through cross-network embedding [48]. This embedding technique enables effective knowledge transfer across drug–target–disease relationships, thereby improving prediction accuracy for both drug-target interactions and drug– disease associations. The iDrug model’s performance was evaluated using a wide range of disease types, demonstrating its broad applicability in repurposing drugs for diverse therapeutic indications. These *in silico* methods open novel strategies for computational drug repurposing. However, the explainability of these methods is still a caveat [49]. The intricate nature of biological systems and the vast amount of data involved in drug repurposing necessitate transparent and interpretable models. In drug repurposing cases, where candidates that have already passed the early stages of drug testing can be directly applied to human subjects, poorly understood or black-box model decisions will arguably not suffice.

In this study we specifically addressed the interpretability by adoption of a data-driven strategy, which shows how a composite FFNN trained on large sets of gene expression data can be used to predict relevant drug repurposing candidates for potential cardiac hypertrophy therapies. We utilized a model structure where groups of known disease-associated genes were compressed independently into one scalar value per disease group. This disease-centered structure was used to predict the full transcriptomic profile, analogously to how regulatory structures were incorporated into the DrugCell by Kuenzi et al. in 2020. The novelty of our approach compared to others is that disease-gene associations, as opposed to cell structures, are directly encoded into the latent space of the model. This strategy provides a more interpretable disease-specific signal in the transcriptome, potentially even offering an alternative to standard gene set annotation analyses, which are often broad and may link to multiple diseases. By investigating the differential response in the hidden nodes of which each corresponds to a specific disease, a direct disease association can be retrieved. Notably, we also show how the model structure successfully captured disease patterns in a wide selection of human diseases. This integration of explainability in the model highlights the potential of neural network learning techniques to encompass a broader understanding of biological mechanisms and pathways, ultimately contributing to the exploration of novel therapeutic avenues.

Leveraging the neural network architecture, we successfully correlated latent disease-associated structures with existing pharmaceutical compounds. We used perturbations of individual nodes as means to estimate the individual node effects on the output layer. While there are some known pitfalls to this approach [50, 51], including the possibility of dependencies between intermediate nodes in relation to model output [52], we were able to predict several potential drug repurposing candidates, notably showcasing lapatinib and amiodarone as promising candidates for mitigating proBNP concentration in cardiac hypertrophy.

ProBNP is a well-known biomarker of cardiac hypertrophy and has elevated levels in patients who suffer from cardiac hypertrophy [53–57]. We therefore used this biomarker for assessing the effect of the candidate drugs. Today, lapatinib is primarily prescribed for breast cancer therapy, whereas amiodarone is typically used to manage different types of arrhythmias [58, 59]. In addition to our candidate drugs, we also included bosentan as a positive control. Bosentan, a dual competitive antagonist of ET-1 receptors, exhibits significantly greater affinity for the ET-A receptor in comparison to the ET-B receptor [60]. The ET-A receptor plays a pivotal role in mediating hypertrophic responses [61]. A plausible explanation for the observed incomplete blunting of ET-1 treatment effects by bosentan lies in the concentration employed during the experiments, which may not have been sufficiently high. In our hypertrophy model, further elevating the concentration was deemed unsuitable due to cell death. An observational study with subjects who are on any of these drugs could be a feasible way to evaluate the clinical implications of these effects. Patients diagnosed with cardiac hypertrophy in such cohort could then be compared with patients in a control cohort.

This study demonstrates the successful development of a straightforward, interpretable modeling approach for efficient identification of existing drugs that can be repurposed for a novel indication. A few top candidates were also experimentally validated in our in vitro disease model of cardiac hypertrophy with significant results. A limitation is the rather few candidate compounds we have tested and validated.

Nevertheless, the scope of this study was to evaluate the applicability of the proposed computational pipeline for identification of candidate drugs for repurposing rather than evaluating novel candidate drugs for cardiac hypertrophy treatment. Future work will involve testing a broader range of compounds and exploring the applicability of the method to other diseases besides cardiac hypertrophy. For those means, a more systematic approach for selecting the most suitable compounds for further *in vitro* validation will be required. However, the aim of this study was to establish the model’s proficiency in predicting gene expressions based on disease-specific variables, which will pave the way for a nuanced understanding of disease states and their molecular underpinnings. Additionally, the model’s interpretable latent space showcased biologically relevant patterns, suggesting a learned comprehension of intricate biological relationships.

Here we used the latent space to embed knowledge associated with diseases, but the technique can be generalized to other biological entities such as molecular functions, biological pathways, or any mechanistic information. The integration of this predictive approach in drug repurposing endeavors could significantly expedite the identification of novel treatments for complex medical conditions like cardiac hypertrophy. This study explores the potential of implementing explainable neural network techniques in drug repurposing, offering insights for future research in predictive medicine and therapeutic exploration.

The future perspective and potential for drug repurposing is amazing and large interest is now reported in using drug repurposing to address the unmet need of therapies for rare diseases with several ongoing global initiatives [62]. The outcomes from these emerging activities will help progress the field with benefits for patients, public health, and medicines development [62–64]. AI-based methods are becoming increasingly important in drug repurposing, but they still face significant challenges with transparency and interpretability, often described as “black-box” issues [34]. While AI has shown great potential for accelerating both drug development and drug repurposing through predictive insights into drug-disease associations, the lack of interpretability remains a significant drawback. More research will be needed to clarify how the AI models reach conclusions and what knowledge that underlies their decisions. This is essential for building trust among researchers and clinicians as well as at the regulatory bodies in the pharmaceutical industry [49]. Here we report on an interpretable FFNN for identification of candidates for drug repurposing, which demonstrates a significant step forward towards explainability and trust in repurposing of drugs.

From the perspective of pharmaceutical companies, the interest in drug repurposing is immense although many and varied challenges remain to be solved [65]. In addition to the scientific challenges of efficient and robust identification of candidate drugs, there is a need to establish business models to support bringing existing molecules as therapies for new indications [65]. Even though the clinical development paths for repurposed drugs may be substantially shortened in drug repurposing, there remains a significant commitment in the need to demonstrate the efficacy and safety of an existing drug in a new indication. Moreover, pharmaceutical companies need to find a way to recoup the investment needed to bring a repurposed product to market. One strategy can be trying to gain exclusivity for the use of a compound for a new indication, which will rely on the generation of new intellectual property [65].

Taken together, successful implementations of drug repurposing approaches have great potential to revolutionize the drug development industry and are anticipated to have a huge impact in many aspects. For patients it will offer faster development of novel therapies with improved safety. The economic benefits are huge, and these resources can instead be redirected to address unmet therapeutic needs for rare disease [62–64]. The need for animal experiments is reduced as pharmacokinetics and pharmacodynamics insights are already known for the repurposed drugs. A more sustainable development of therapeutic avenues is gained by offering a pathway to bring treatments to market faster and at a lower cost with increased access to affordable treatments for rare diseases that might otherwise lack effective therapies.

## 5. Conclusions

This study demonstrated how a composite feed-forward neural network can predict relevant drug repurposing candidates for treatment of cardiac hypertrophy, and validated their performance in an in vitro cardiac hypertrophy model. Specifically, we used a FFNN structure with an explainable latent space that was tailored to correspond to disease signals in gene expression data. This model was used to extract a disease vector based on RNA-Seq data of ET-1 stimulated hiPSC-derived CMs represented in the latent space, and to connect this vector to known drug targets. Based on these candidate predictions, we experimentally validated the effects of lapatinib and amiodarone in our hypertrophy model based on hiPSC-derived cardiomyocytes. In summary, although the number of drugs that we have validated is limited, this work demonstrates an interesting approach where AI techniques are utilized for drug repurposing in a way that facilitates biological interpretation of the learned embeddings, shedding light on its immense promise in reshaping the landscape of drug discovery and development.

## Acknowledgement

Figure 1, 2 and 6 were created with the BioRender software (www.biorender.com).

## Data Availability

RNA-seq data from the cardiac hypertrophy model can be accessed at: https://www.ebi.ac.uk/biostudies/arrayexpress/studies/E-MTAB-11030

## Supplementary Figure legends

**Supp. Fig 1.** proBNP concentration analysis of all substances tested. Initial screening to investigate which substances that reduced the proBNP concentration. The y-axis represents the concentration in ng/ml, and x-axis shows the different treatment groups. The low proBNP concentration in Regorafinib (10 µM) is attributed to cell death.

**Supp. Fig. 2.** Brightfield images captured immediately before sample collection. Magnification for all images is 10x. A) Amiodarone 1.0uM B) Amiodarone 10uM C) Lapatinib 0.1uM D) Lapatinib 1.0uM E) Lapatinib 10uM F) Bosentan 10uM G) Bosentan 100uM

**Suppl. Fig. 3.** Brightfield images captured immediately before sample collection. Magnification for all images is 10x. A) Control (DMSO) B) Control (Methanol) C) ET-1 (DMSO) D) ET-1 (Methanol)

---

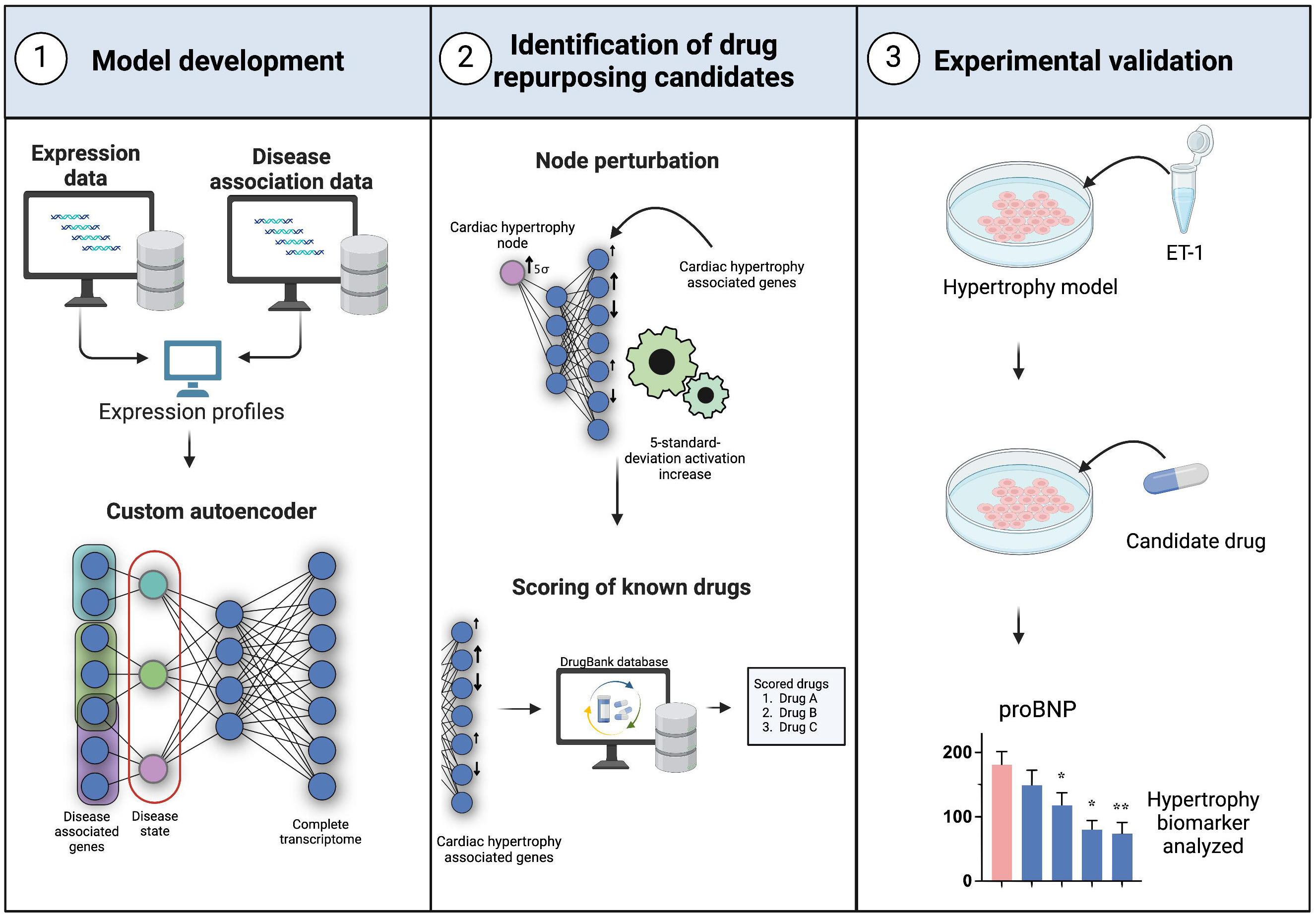

## References

1. Verbaanderd, C., I. Rooman, and I. Huys, Exploring new uses for existing drugs: innovative mechanisms to fund independent clinical research. Trials, 2021. 22(1): p. 322.

2. Luca, P., B. Nicolo, and R. Giulio, How drug repurposing can advance drug discovery: challenges and opportunities. Frontiers in Drug Discovery, 2014. 4.

3. Ashburn, T.T. and K.B. Thor, Drug repositioning: identifying and developing new uses for existing drugs. Nat Rev Drug Discov, 2004. 3(8): p. 673–83.

4. Corsello, S.M., et al., The Drug Repurposing Hub: a next-generation drug library and information resource. Nat Med, 2017. 23(4): p. 405–408.

5. Oprea, T.I. and J. Mestres, Drug repurposing: far beyond new targets for old drugs. AAPS J, 2012. 14(4): p. 759–63.

6. Pushpakom, S., et al., Drug repurposing: progress, challenges and recommendations. Nat Rev Drug Discov, 2019. 18(1): p. 41–58.

7. Chung, C.C.Y., et al., Rare disease emerging as a global public health priority. Frontiers in Public Health Policy, 2022. 10.

8. Ghandikota, S.K. and A.G. Jegga, Application of artificial intelligence and machine learning in drug repurposing. Prog Mol Biol Transl Sci, 2024. 205: p. 171–211.

9. Troseid, M., et al., Efficacy and safety of baricitinib in hospitalized adults with severe or critical COVID-19 (Bari-SolidAct): a randomised, double-blind, placebo-controlled phase 3 trial. Crit Care, 2023. 27(1): p. 9.

10. Parvathaneni, V., et al., Drug repurposing: a promising tool to accelerate the drug discovery process. Drug Discov Today, 2019. 24(10): p. 2076–2085.

11. Jin, G. and S.T. Wong, Toward better drug repositioning: prioritizing and integrating existing methods into efficient pipelines. Drug Discov Today, 2014. 19(5): p. 637–44.

12. Peyvandipour, A., et al., A novel computational approach for drug repurposing using systems biology. Bioinformatics, 2018. 34(16): p. 2817–2825.

13. Saberian, N., et al., A new computational drug repurposing method using established disease-drug pair knowledge. Bioinformatics, 2019. 35(19): p. 3672–3678.

14. Karatzas, E., G. Kolios, and G.M. Spyrou, An Application of Computational Drug Repurposing Based on Transcriptomic Signatures. Methods Mol Biol, 2019. 1903: p. 149–177.

15. Chen, J., et al., Deep transfer learning of cancer drug responses by integrating bulk and single-cell RNA-seq data. Nat Commun, 2022. 13(1): p. 6494.

16. Kuenzi, B.M., et al., Predicting Drug Response and Synergy Using a Deep Learning Model of Human Cancer Cells. Cancer Cell, 2020. 38(5): p. 672–684 e6.

17. Pinero, J., et al., The DisGeNET knowledge platform for disease genomics: 2019 update. Nucleic Acids Res, 2020. 48(D1): p. D845–D855.

18. Frey, N., et al., Hypertrophy of the heart: a new therapeutic target? Circulation, 2004. 109(13): p. 1580–9.

19. Moreira, J.B.N., M. Wohlwend, and U. Wisloff, Exercise and cardiac health: physiological and molecular insights. Nat Metab, 2020. 2(9): p. 829–839.

20. Eghbali, M., et al., Molecular and functional signature of heart hypertrophy during pregnancy. Circ Res, 2005. 96(11): p. 1208–16.

21. Tham, Y.K., et al., Pathophysiology of cardiac hypertrophy and heart failure: signaling pathways and novel therapeutic targets. Arch Toxicol, 2015. 89(9): p. 1401–38.

22. Hieda, M., et al., Increased Myocardial Stiffness in Patients With High-Risk Left Ventricular Hypertrophy: The Hallmark of Stage-B Heart Failure With Preserved Ejection Fraction. Circulation, 2020. 141(2): p. 115–123.

23. Frey, N. and E.N. Olson, Cardiac hypertrophy: the good, the bad, and the ugly. Annu Rev Physiol, 2003. 65: p. 45–79.

24. Hill, J.A. and E.N. Olson, Cardiac plasticity. N Engl J Med, 2008. 358(13): p. 1370–80.

25. Nakamura, M. and J. Sadoshima, Mechanisms of physiological and pathological cardiac hypertrophy. Nat Rev Cardiol, 2018. 15(7): p. 387–407.

26. Oka, T. and I. Komuro, Molecular mechanisms underlying the transition of cardiac hypertrophy to heart failure. Circ J, 2008. 72 **Suppl A**: p. A13-6.

27. Martin, T.G., M.A. Juarros, and L.A. Leinwand, Regression of cardiac hypertrophy in health and disease: mechanisms and therapeutic potential. Nat Rev Cardiol, 2023. 20(5): p. 347–363.

28. Lachmann, A., et al., Massive mining of publicly available RNA-seq data from human and mouse. Nat Commun, 2018. 9(1): p. 1366.

29. de Weerd, H.A., et al., Representational Learning from Healthy Multi-Tissue Human RNA-seq Data such that Latent Space Arithmetics Extracts Disease Modules. https://www.biorxiv.org/content/10.1101/2023.10.03.560661v1, 2024.

30. Kingma, D.P. and J. Ba, Adam: A Method for Stochastic Optimization. arXiv:1412.6980v9 [cs.LG], 2017.

31. Johansson, M., et al., Multi-Omics Characterization of a Human Stem Cell-Based Model of Cardiac Hypertrophy. Life (Basel), 2022. 12(2).

32. Guney, E., et al., Network-based in silico drug efficacy screening. Nat Commun, 2016. 7: p. 10331.

33. Dwivedi, S.K., et al., Deriving disease modules from the compressed transcriptional space embedded in a deep autoencoder. Nat Commun, 2020. 11(1): p. 856.

34. Magnusson, R., J.N. Tegner, and M. Gustafsson, Deep neural network prediction of genome-wide transcriptome signatures - beyond the Black-box. NPJ Syst Biol Appl, 2022. 8(1): p. 9.

35. Johansson, M., et al., Cardiac hypertrophy in a dish: a human stem cell based model. Biol Open, 2020. 9(9).

36. Hink, U., et al., Role for peroxynitrite in the inhibition of prostacyclin synthase in nitrate tolerance. J Am Coll Cardiol, 2003. 42(10): p. 1826–34.

37. Pratt, C.M., et al., Mortality in the Survival With ORal D-sotalol (SWORD) trial: why did patients die? Am J Cardiol, 1998. 81(7): p. 869–76.

38. Johansson, M., et al., Data Mining Identifies CCN2 and THBS1 as Biomarker Candidates for Cardiac Hypertrophy. Life (Basel), 2022. 12(5).

39. Mullard, A., 2019 FDA drug approvals. Nat Rev Drug Discov, 2020. 19(2): p. 79–84.

40. Pammolli, F., L. Magazzini, and M. Riccaboni, The productivity crisis in pharmaceutical R&D. Nat Rev Drug Discov, 2011. 10(6): p. 428–38.

41. Kaitin, K.I., Deconstructing the drug development process: the new face of innovation. Clin Pharmacol Ther, 2010. 87(3): p. 356–61.

42. DiMasi, J.A., H.G. Grabowski, and R.W. Hansen, Innovation in the pharmaceutical industry: New estimates of R&D costs. J Health Econ, 2016. 47: p. 20–33.

43. A, N., O. N, and D.D. L, Renovation as innovation: is repurposing the future of drug discovery research? Drug Discovery today, 2018. 24.

44. C, G. and M. A, Is drug repurposing really the future of drug discovery or is new innovation truly the way forward?. Expert Opinion on Drug Discovery, 2021. 16.

45. Kumar, S. and V. Roy, Repurposing Drugs: An Empowering Approach to Drug Discovery and Development. Drug Res (Stuttg), 2023. 73(9): p. 481–490.

46. Schlander, M., et al., How Much Does It Cost to Research and Develop a New Drug? A Systematic Review and Assessment. Pharmacoeconomics, 2021. 39(11): p. 1243–1269.

47. Zhao, T., et al., Identifying drug-target interactions based on graph convolutional network and deep neural network. Brief Bioinform, 2021. 22(2): p. 2141–2150.

48. Chen, H., F. Cheng, and J. Li, iDrug: Integration of drug repositioning and drug-target prediction via cross-network embedding. PLoS Comput Biol, 2020. 16(7): p. e1008040.

49. Alizadehsani, R., et al., Explainable Artificial Intelligence for Drug Discovery and Development: A Comprehensive Survey. IEEE Access, 2024. 12.

50. A, d.W., et al., Latent space arithmetic on data embeddings from healthy multi-tissue human RNA-seq decodes disease modules. Patterns, 2024.

51. Kingma, D.P. and M. Welling, Auto-Encoding Variational Bayes, in In 2nd International Conference on Learning Representations (ICLR). 2014.

52. Ribeiro, M.T., S. Singh, and C. Guestrin, “Why Should I Trust You?” Explaining the Predictions of Any Classifier. Bioxarchive, 2016. https://arxiv.org/pdf/1602.04938.pdf.

53. Coats, C.J., et al., Cardiac biomarkers and effects of aficamten in obstructive hypertrophic cardiomyopathy: the SEQUOIA-HCM trial. Eur Heart J, 2024.

54. Coats, C.J., et al., Relation between serum N-terminal pro-brain natriuretic peptide and prognosis in patients with hypertrophic cardiomyopathy. Eur Heart J, 2013. 34(32): p. 2529–37.

55. Rivera Otero, J.M., et al., Ventricular hypertrophy increases NT-proBNP in subjects with and without hypertension. Int J Cardiol, 2004. 96(2): p. 265–71.

56. Morillas, P., et al., Usefulness of NT-proBNP level for diagnosing left ventricular hypertrophy in hypertensive patients. A cardiac magnetic resonance study. Rev Esp Cardiol, 2008. 61(9): p. 972–5.

57. Sergeeva, I.A. and V.M. Christoffels, Regulation of expression of atrial and brain natriuretic peptide, biomarkers for heart development and disease. Biochim Biophys Acta, 2013. 1832(12): p. 2403–13.

58. Hamilton, D., Sr., et al., Amiodarone: A Comprehensive Guide for Clinicians. Am J Cardiovasc Drugs, 2020. 20(6): p. 549–558.

59. Voigtlaender, M., T. Schneider-Merck, and M. Trepel, Lapatinib. Recent Results Cancer Res, 2018. 211: p. 19–44.

60. Martin, C., H.D. Held, and S. Uhlig, Differential effects of the mixed ET(A)/ET(B)-receptor antagonist bosentan on endothelin-induced bronchoconstriction, vasoconstriction and prostacyclin release. Naunyn Schmiedebergs Arch Pharmacol, 2000. 362(2): p. 128–36.

61. Marasciulo, F.L., M. Montagnani, and M.A. Potenza, Endothelin-1: the yin and yang on vascular function. Curr Med Chem, 2006. 13(14): p. 1655–65.

62. Jonker, A.H., et al., Drug repurposing for rare: progress and opportunities for the rare disease community. Front Med (Lausanne), 2024. 11: p. 1352803.

63. Cortial, L., et al., Artificial intelligence in drug repurposing for rare diseases: a mini-review. Front Med (Lausanne), 2024. 11: p. 1404338.

64. Fetro, C. and D. Scherman, Drug repurposing in rare diseases: Myths and reality. Therapie, 2020. 75(2): p. 157–160.

65. Cha, Y., et al., Drug repurposing from the perspective of pharmaceutical companies. Br J Pharmacol, 2018. 175(2): p. 168–180.

