## Supplementary figures and images for "Leveraging Neural Network Models for Drug Repurposing: A case study on Cardiac Hypertrophy"

### Supplementary figure 1

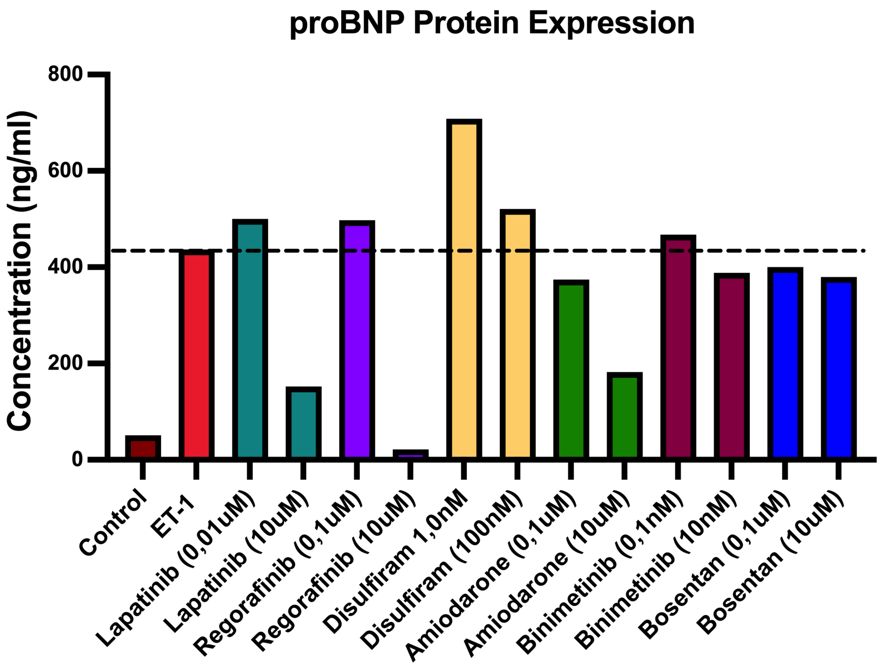

### Supplementary figure 2

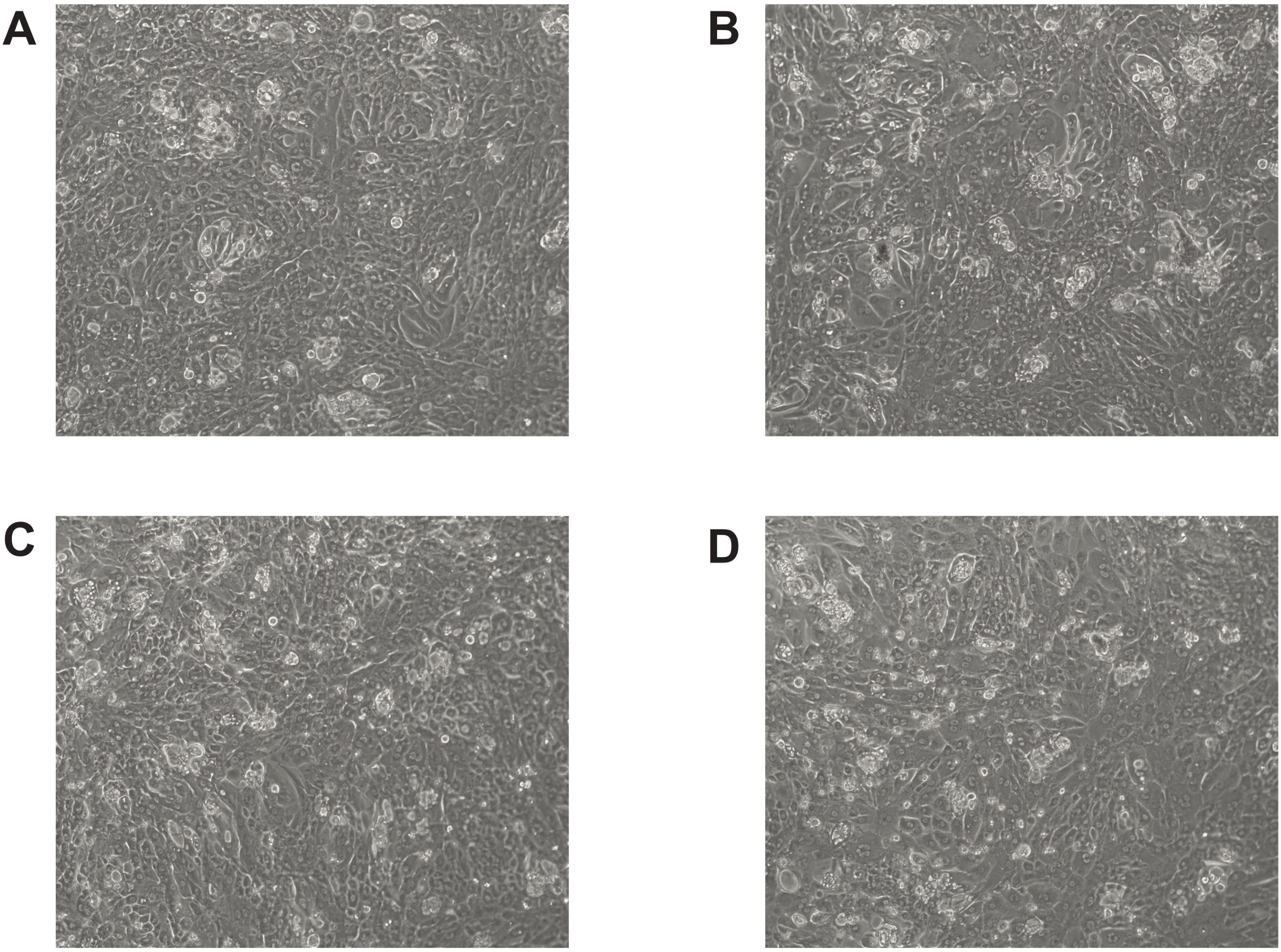

### Supplementary figure 3

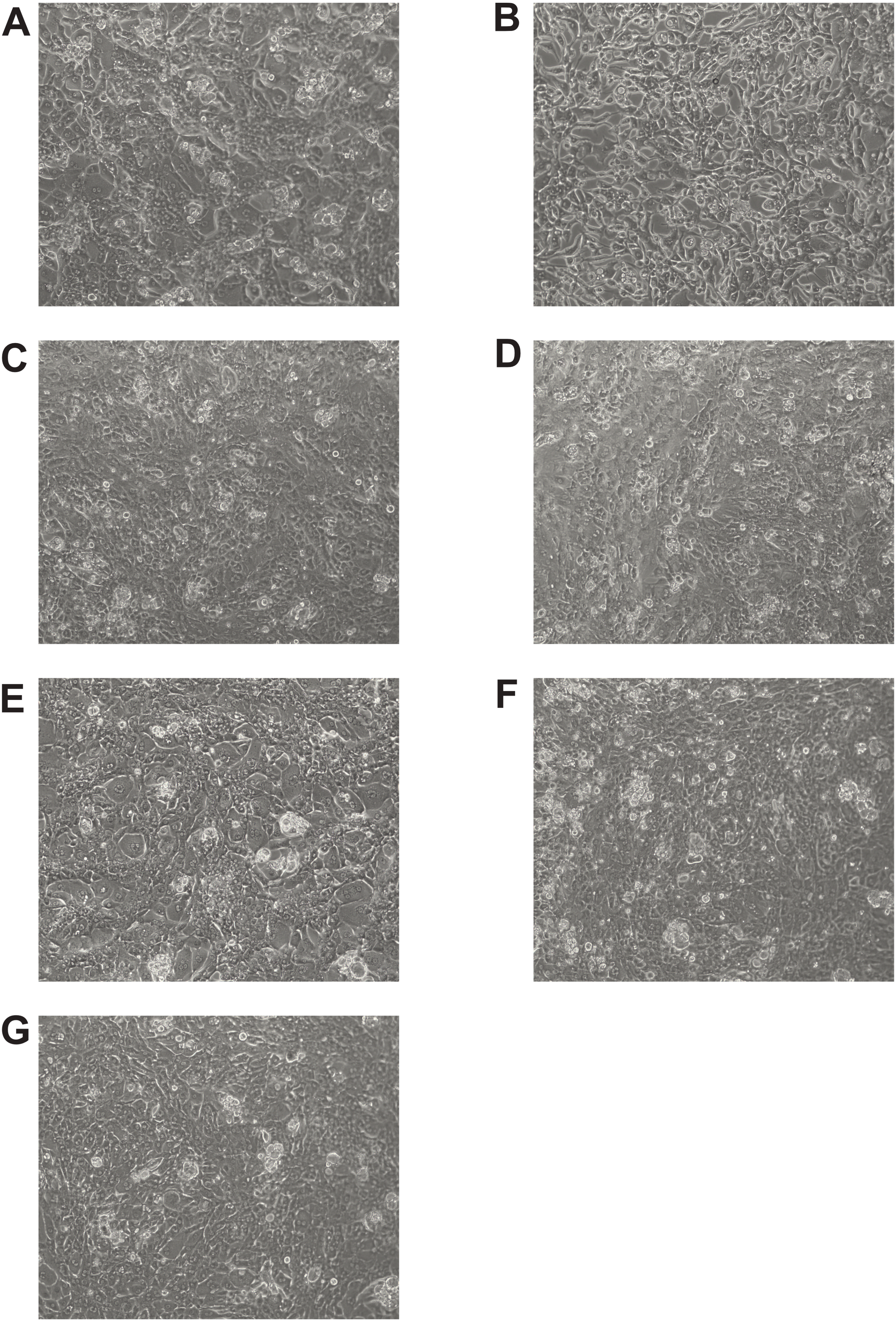
